# Separation of Bioactive Small Molecules, Peptide from Natural Products and Proteins from Pathogenic Microbe by Rotofor-based Preparative Isoelectric Focusing (IEF) Method

**DOI:** 10.1101/2019.12.23.887711

**Authors:** Raja Veerapandian, Anuja Paudyal, Adeline Chang, Govindsamy Vediyappan

## Abstract

The primary purpose of purifying the bioactive small molecules is obtaining enough amount of specific molecules for their downstream applications. There are various methods available for serving this purpose. However, not all can afford for cost-effective purification methods, especially for complex biological samples. Here we demonstrate, the Rotofor based preparative IEF separation which shows the high possibility of separating and enriching desired molecules, including small molecules and peptides from the complex plant extract. This method is well-suited for use at any stage of a purification scheme and has been used for complex biological samples quantification and characterization. As a proof of concept, we fractionated *Gymnema sylvestre* plant extract and isolated a family of terpenoid saponin molecules at the acidic end and small peptide molecule near the basic end. To our knowledge, this is a first report that Rotofor can be used to fractionate bioactive small molecules from natural product sources. We also showed effective microbial protein separation using *Candida albicans* fungus as a model system. The Rotofor based IEF has appealing features such as low cost, simplicity, high application potential and can be performed in any laboratory with the instrument setup.

## Introduction

The purification of biomolecules is the essential and difficult task of any biological experiments, especially for a complex biological sample^1^. Biomolecules purification shows numerous challenges due to their high complexity, low abundance, and extremes in their pI values and other physicochemical properties^2^. Isoelectric focusing (IEF) is one of the finest techniques with high resolution separation of complex biomolecules where ampholytes travel according to their charge in the established pH gradient under the influence of an electric field until the net charge of the molecule is zero^3^. There are several methods used for separation of biomolecules like agarose and polyacrylamide gel electrophoresis, two-dimensional gel electrophoresis (2-DE), capillary electrophoresis, isotachophoresis and various chromatographic techniques (TLC, FPLC, HPLC, etc.)^2^.

Rotofor (Bio-Rad, Hercules, CA, USA), an IEF based apparatus consist of cylindrical focusing chambers divided into 20 compartments and it provides up to 500-fold purification for a specific molecule in < 4 hours. The most advantageous part of Rotofor is that it can be applied at any stage of the purification process. For example, it can be used to purify complex biological samples like animal tissue or plant leaf extract or for purifying the few nonspecific contaminants in the pre purified protein sample^4^. Also, during the entire process the protein native conformation is maintained as the experiment is run entirely in a free solution with controlled temperature.

Even though the use of Rotofor is reported widely for proteins, enzymes and antibody purifications^4–8^, here we describe the use of Rotofor for separating and purifying small molecules and peptides from the medicinal plant called *Gymnema sylvestre*. This will help the researchers to concentrate and purify their active small molecules from a mixture of plant extract for their downstream applications with less cost under native conditions. In addition, we also show the enrichment of certain proteins from a complex protein extract from *Candida albicans* fungus^9^ in Rotofor system as an example.

## Protocol

### 1. Setup and prerunning of standard Rotofor unit

1. Equilibration of ion exchange membranes: The electrodes (anode -red button and cathode - black button) with its respective exchange membrane were assembled according the manufacturer’s instructions. The anion exchange membranes were equilibrated in 0.1 M NaOH and the cation exchange membranes were equilibrated with 0.1 M H_3_PO_4_. **Note:** The membranes should be stored in electrolytes between runs and not to be dried out.
2. Electrode assembly: The inner and outer portion of the electrode were assembled by aligning three oblong holes in the ion-exchange gaskets. Fill the electrodes with respective electrolytes (~25-30 mL) to prevent membrane from drying. **Note:** Don’t add excess electrolyte which may build up pressure inside the electrode and cause leaking.
3. Focusing chamber assembly: All the parts (anode electrode, membrane core, focusing chamber and cathode electrode) were assembled over the ceramic cooling finger. Cover the sample collection ports with a sealing tape. **Note:** Refer Bio-Rad’s Rotofor® system instruction manual for detailed assembly instructions.
4. Connect the Rotofor to the coolant with 4ºC and prerun the unit with distilled water until the voltage stabilizes. **Note:** Prerun with distilled water helps to remove the residual base from the focusing chamber and the nylon membrane core.

### 2. Separation and purification of small molecules and peptides from *Gymnema sylvestre* plant extract

1. The plant extract (25% methanol extract, Suan Forma, NJ, USA) of *G. sylvestre* were weighed (0.6 g) and dissolved in distilled water (60 mL) and mixed well. **Note:** Any complex biological samples that are soluble and free of salt can be used for the separation and purification using Rotofor. We have studied the saponin family of triterpenoid compounds called gymnemic acids from *G. sylvestre* plant for their unique antifungal properties^10^. Any plant extract with above mentioned properties can be used in Rotofor based IEF.
2. Then the solubilize plant extract was briefly centrifuged at 10,000 x g for 5 min to remove insoluble particles.
3. The supernatant was transferred to the new tube and mixed with ampholyte (pH 3-10, Bio-rad). We employed 1% (v/v) ampholyte concentration. **Note:** An ampholyte concentration of up to 2% can be used. Always keep the samples, ampholyte and water precooled on the ice.
4. Follow the steps from protocol no 1. Setup Rotofor assembly and prerun the standard Rotofor unit for 3-5 minutes at 15W, 3000 Voltage or as per the Rotofor manual guidelines. Generally, within one minute, the voltage will reach the maximum set values. Then switch off the power source and remove the water from the cell using the fraction collector. Re-seal the collection ports with sealing tape. Now the chamber is ready for loading the sample.
5. Load the prepared sample with ampholyte in the cell through sample injection ports. A 50 ml syringe with a 1-1/2 inch 19-gauge needle was used for sample loading.
6. Remove air bubbles from the sample cell. The presence of air bubbles will cause voltage and current fluctuations and affect the run. **Note:** To remove air bubbles, remove the sample cell from the stand and tap the electrode chamber to dislodge the bubbles.
7. Connect the unit to the coolant (4ºC) and start fractionation with the constant power supply. We achieved a maximum of 750 voltage, 20 mA out of the set parameters (15W, 300 mA & 3000 voltage) in 4 hours of focusing at 5°C circulating water temperature.
8. Run the apparatus for 3 h or until the voltage reached a constant value.
9. After the run, fractions were collected in the harvest box (contains 20 tubes) connected to the vacuum pump by pressing the harvest button “ON”.
10. Fractions were used for subsequent downstream applications (SDS-PAGE for peptide quantification, TLC and bioassay for small molecules).

### 3. Separation and purification of proteins from *C. albicans*

1. Cell surface protein extract (non-glucan attached cell wall proteins) from *C. albicans* yeast cells was prepared with ammonium carbonate (1.89 g/L) buffer containing 1% (v/v) beta-mercaptoethanol (βME) (1/10^th^ of the culture volume) as described^9^. **Note:** Protein samples from *C. albicans* cytosol, membrane or cell wall can be extracted by different methods and can be used for Rotofor fractionation. Similarly, proteins from bacteria or other biological samples (animal tissue extract or washes) can be used after removing any salt by appropriate methods.
2. The protein extract was filtered (0.2 μM filter) and dialyzed against water. The protein concentration was estimated by the Bradford dye-binding method using gamma globulin as a standard.
3. About 500 mg of total protein in 60 ml of water containing 1% (v/v) ampholyte (pH 5-8) was used to fractionate the proteins. The maximum voltage of 2020 and 10 mA out of the set parameters (15W, 300 mA & 3000 voltage) for a total of 4 hours focusing at 5°C circulating water temperature were achieved. **Note:** Since the broad-range ampholyte (pH 3-10) doesn’t enrich certain non-glucan attached cell wall proteins of *C. albicans* well, we used a narrow-range (pH 5-8) ampholyte.
4. After 4 hours of focusing, protein fractions (1-20) were harvested as described above and analyzed on 12.5% SDS-PAGE after reducing and boiling the protein samples.
5. Resolved proteins were stained with Coomassie blue dye and the gel was photographed.

### 4. Bioactivity of purified small molecules from plant extract *Gymnema sylvestre*

1. Overnight grown *C. albicans* yeast cells were diluted (1/1000 dilution) into a fresh RPMI cell culture medium supplemented with 50 mM glucose. **Note:** *C. albicans* converts from yeast growth to hyphae under hyphal inducing conditions (RPMI at 37°C). Gymnemic acid small molecules are shown to inhibit the conversion of yeast cells into hyphae under hypha inducing conditions^10^. We aim to determine if Rotofor separated *G. sylvestre* extract contain these bioactive molecules.
2. From this cell suspension, 90 μL was added to each well of the 96-well plate.
3. From each fraction, 10 μL (1-20 harvested fractions, Figure 4) of *G. sylvestre* extract obtained from Rotofor IEF were added into the wells. The assay was performed in triplicates.
4. As a negative control, 10 μL of ampholyte in water (1%) was also used in separate wells. Note: It is known that ampholytes have bioactivity potential and so it is essential to include ampholytes control while performing any bioassay^11^.
5. The 96-well plate was incubated at 37°C for 12 hours and inhibition of *C. albicans* yeast to hypha conversion was observed under the microscope.

## Results and Discussion

Small molecules from natural product sources (*e.g.* plants) are complex secondary metabolites that are highly diverse in chemical structures and are believed to be involved in plant defense mechanisms. Similarly, polypeptides are known to be present in plant tissues ^12^. These natural product small molecules are rich sources for drug discovery and development. However, the difficulties and tedious methods required for their isolation and purification limit their use for therapeutic applications. The Rotofor IEF approach we used in this report highlights the ease of separating these small molecules and polypeptides without compromising their bioactivities.

Using Rotofor preparative IEF method, we have fractionated a medicinal plant extract and cell surface proteins from a human pathogenic fungus, *C. albicans*. A schematic of these fractionation protocols is shown in Figure 1.

**Figure 1:**
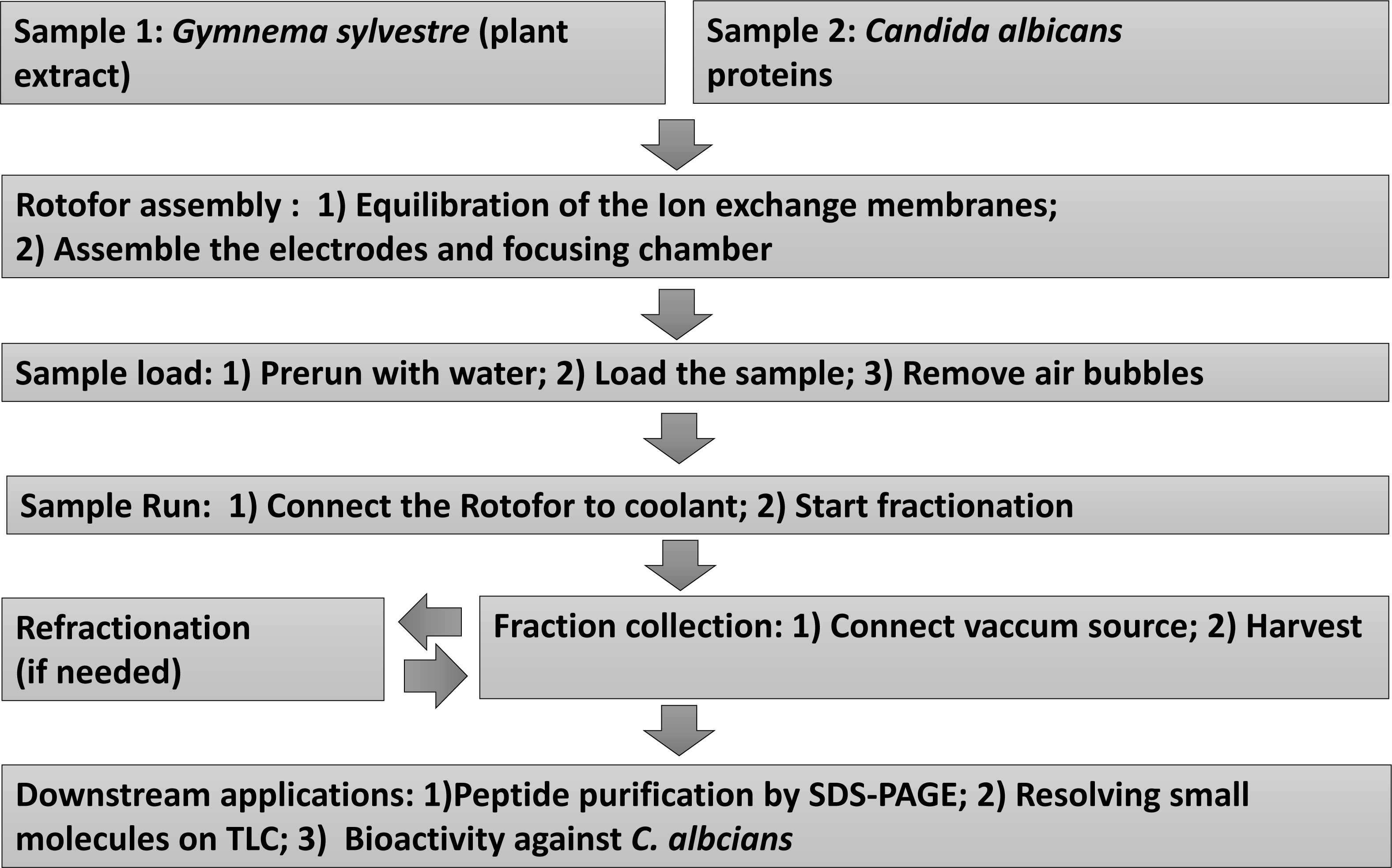
Flow chart showing the experimental work flow. Stepwise Rotofor fractionation procedures and subsequent downstream assays are mentioned. Samples include *Gymnema sylvestre* (leaf extract, sample 1) and *Candida albicans* non-glucan attached yeast proteins (sample 1).

Out of 20 fractions of *G. sylvestre* extract obtained from Rotofor IEF, the dark-colored molecules (terpenoid saponins) were migrated and enriched at the anode end (pH 2-3) and the light-yellow clear fractions were observed at cathode end (pH 8-9) (Figure 2). Aliquots (20 μl) from each fraction (1-20) were separated after reducing and boiling on 15% SDS-PAGE. The Coomassie blue-stained gel shows the diffused polypeptide band of about 5 kDa that is enriched in fractions 16-19 (Figure 3). It has been reported that *G. sylvestre* plant contains a 35 amino acid gurmarin basic polypeptide with the predicted molecular weight of 4,209^13^. This polypeptide is known to suppress sweet-taste sensation^13^. Bacteria, plants and animals contain peptides, many of them are circular (knottins) and stable with wide range of biological activities such as insecticidal and antimicrobial^12,14^.

**Figure 2:**
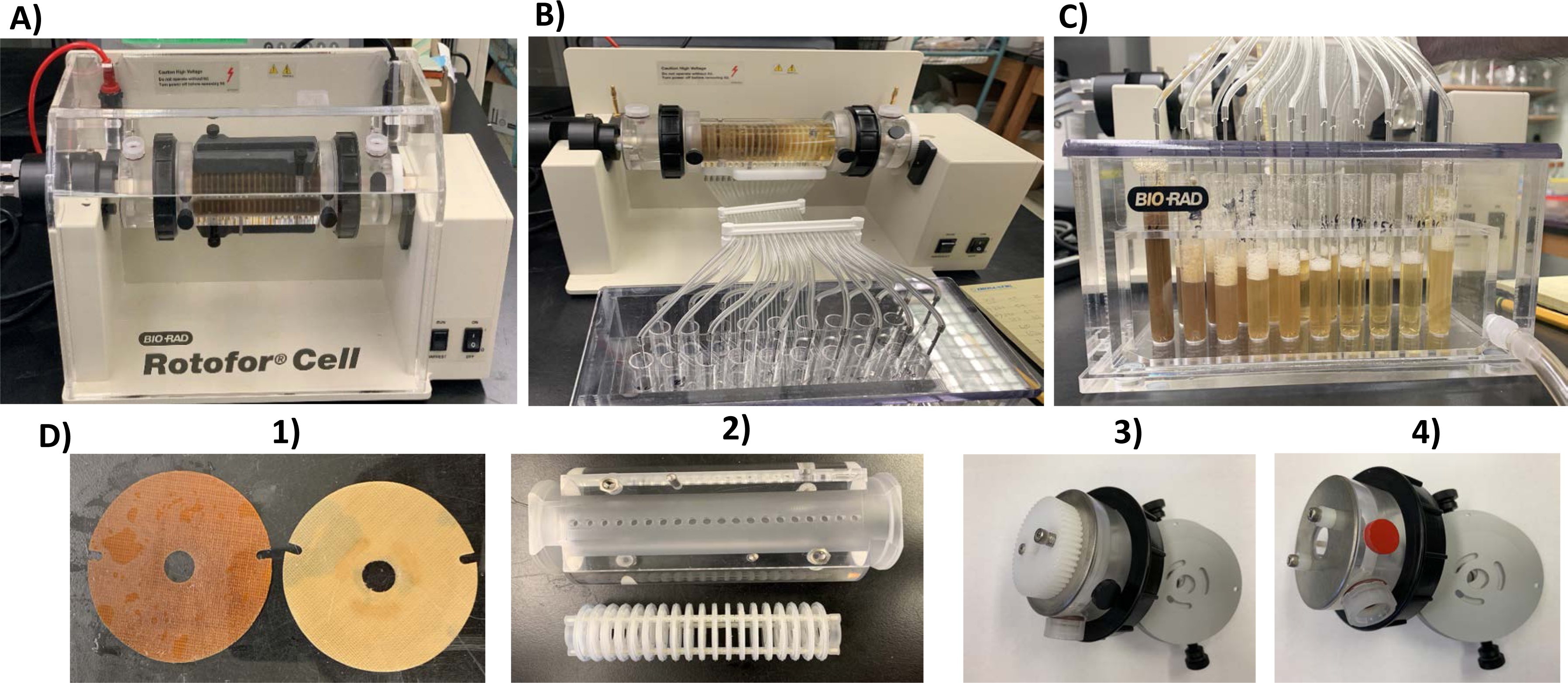
Picture showing the Rotofor apparatus setup and fractionation of *G. sylvestre* plant extract. **A**) During Run, **B**) During fraction collection, **C**) After fraction collection, **D**) Rotofor parts, 1) Ion exchange membranes, 2-Focusing chamber and membrane core, 3-Electrode assembly (negative), 4-Electrode assembly (positive).

**Figure 3:**
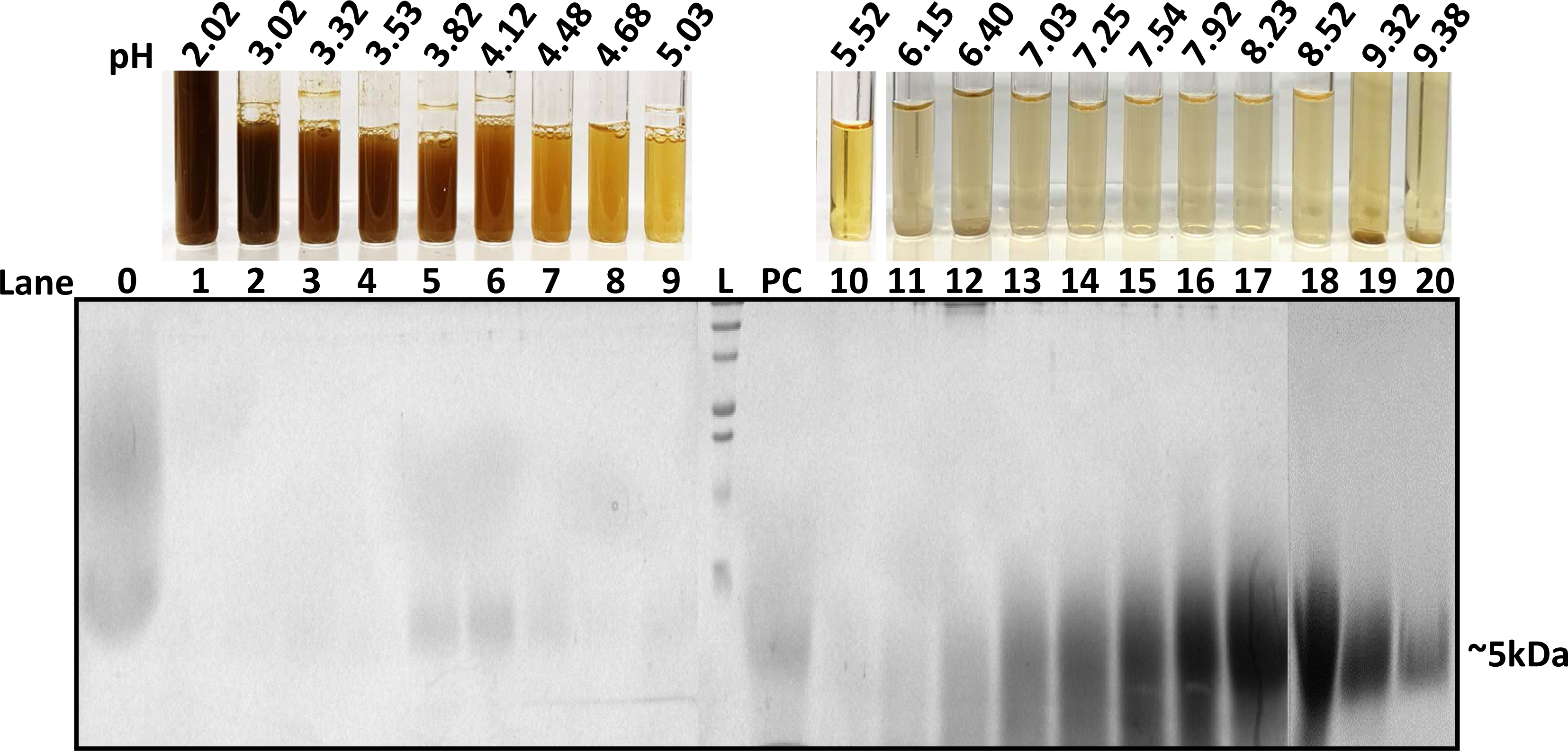
SDS-PAGE separations of the Rotofor plant extract fractions showing peptide. L-ladder, PC-Positive control (peptide), 0 - input sample, 1-20 separated Rotofor fractions. SDS-PAGE (15% resolving gel) was stained by Coomassie blue dye to visualize the resolved peptides (~5 kDa) from fractions 1-20. Fractions 1-3 contain small molecules (fraction 1 has darker color indicating enriched compounds) which can’t be stained/detected by Coomassie blue dye.

*G. sylvestre* plant also contains gymnemic acids (terpenoid saponins) as major constituents^10,15,16^. As expected, these small molecules in fraction 1 and few next fractions were not detected by SDS-PAGE and Coomassie staining (Figure 3). However, these small molecules can be separated by TLC and detected under the UV light (Figure 4, panel A, lane F1). Fraction 10 did not contain a detectable amount of these small molecules on TLC suggesting most of the organic small molecules were enriched in fractions 1-3. Gymnemic acids (GAs) molecules were shown to inhibit *C. albicans* yeast to hypha transition^10,17^. We determined all the 20 fractions collected in this study for their inhibitory activity against *C. albicans* yeast to hypha conversion and hyphal growth. The results are shown in Figure 4, panel B & C. The highest activity is observed in fraction 1 which agrees with the TLC results where several spots can be seen. Isomers of gymnemic acids exist and all of them have similar biological activities^10^. These isomers were separated in fraction 1-3 and show inhibition of *C. albicans* hyphal growth (panels A & C, fraction 1). The degree of hyphal inhibition was gradually decreased as it goes from 1 to 10. Least or no activity was obtained in fractions 10 and above.

**Figure 4:**
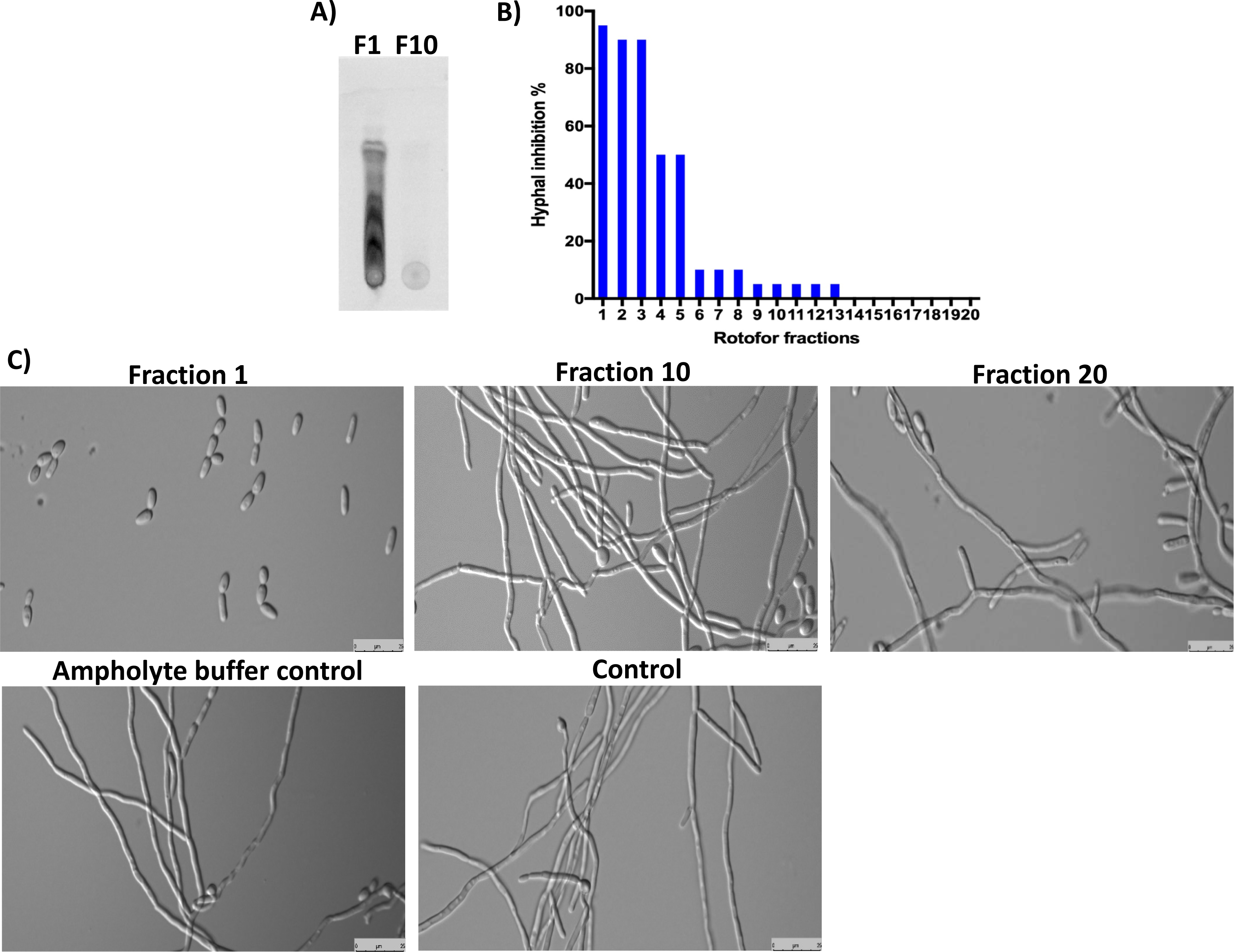
Analysis of Rotofor fractionated small molecules on TLC and determination of their bioactivity against *C. albicans*. Panel **A**) shows TLC analysis of small molecules from fraction #1 and #10. Activated silica gel plate was used to spot ~5 µl of samples and ran with toluene: chloroform: methanol solvent (5:8:3 ratio) until the solvent front reached the margin. TLC separated compounds were detected under an epifluorescence UV light (310 nm). Panel **B**) shows the % inhibition of *C. albicans* yeast to hypha conversion by different fractions. Panel **C**) demonstrates the cell morphology of *C. albicans* under hypha inducing conditions. Fraction #1 shows maximum (98%) inhibition of yeast to hypha conversion. Other fractions and controls show no inhibition of *C. albicans* hyphal growth after 12 hours of incubation at 37°C.

Results from Rotofor fractionation of *C. albicans* cell surface proteins are shown in Figure 5. These cell surface proteins play important roles in *C. albicans* adhesion and pathogenesis. Several enriched proteins (arrows) in different fractions were observed which may be useful to identify for their immunological reactions with Candida infected human serum and identify by mass spectrometry. The Rotofor based purification plays a crucial role in biomolecules separation and purification, thus, when coupled with mass spectrometry analysis it will allow us to identify low abundance proteins from complex biological samples.

**Figure 5:**
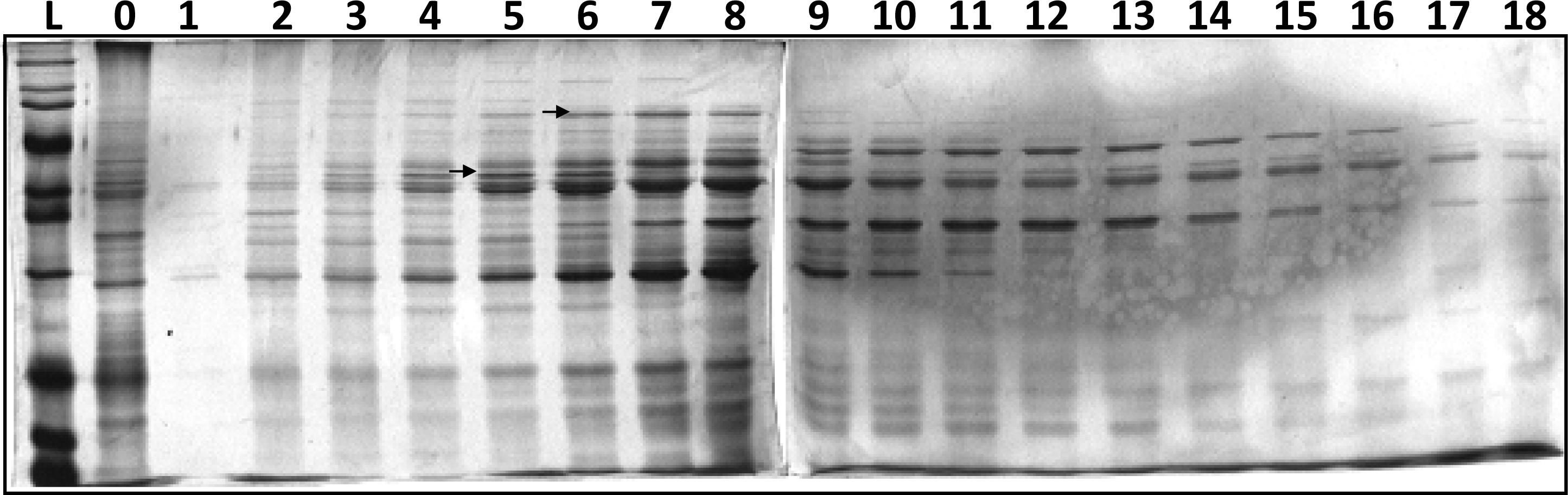
SDS-PAGE analysis of *C. albicans* cell surface proteins (non-glucan attached). L-ladder, 0 - input sample, 1-18 fractions collected after IEF in a standard Rotofor cell using a narrow range (pH 5-8) ampholyte. The image shows SDS-PAGE (12.5%) resolved proteins after staining with Coomassie blue dye. Several proteins were enriched in certain fractions (arrows).

In summary, using the Rotofor IEF method we have shown the separation of bioactive gymnemic acids and gurmarin polypeptide from *G. sylvestre* leaf extract. To our knowledge, we are the first one to register that Rotofor can also be used to purify bioactive small molecules from natural product sources apart from proteins and peptides purification. Further, this IEF method can be useful to enrich selective proteins from the complex crude extracts of pathogenic microbes.

## Acknowledgments

We are thankful for the funding sources from the Division of Biology and Johnson Cancer Research Center for BRIEF and IRA awards, respectively to GV. We also thank the K-INBRE postdoctoral award to RV. This work was supported in part by the Institutional Development Award (IDeA) from the National Institute of General Medical Sciences of the National Institutes of Health under grant number P20 GM103418. The content is solely the responsibility of the authors and does not necessarily represent the official views of the National Institute of General Medical Sciences or the National Institutes of Health.

